# Genetic variation across the human olfactory receptor repertoire alters odor perception

**DOI:** 10.1101/212431

**Authors:** Casey Trimmer, Andreas Keller, Nicolle R. Murphy, Lindsey L. Snyder, Jason R. Willer, Maira Nagai, Nicholas Katsanis, Leslie B. Vosshall, Hiroaki Matsunami, Joel D. Mainland

## Abstract

The human olfactory receptor repertoire is characterized by an abundance of genetic variation that affects receptor response, but the perceptual effects of this variation are unclear. To address this issue, we sequenced the OR repertoire in 332 individuals and examined the relationship between genetic variation and 276 olfactory phenotypes, including the perceived intensity and pleasantness of 68 odorants at two concentrations, detection thresholds of three odorants, and general olfactory acuity. Genetic variation in a single OR frequently associated with odorant perception, and we validated 10 cases in which in vitro OR function correlated with in vivo odorant perception using a functional assay. This more than doubles the published examples of this phenomenon. For eight of these 10 cases, reduced receptor function associated with reduced intensity perception. In addition, we used participant genotypes to quantify genetic ancestry and found that, in combination with single OR genotype, age and gender, we can explain between 10 and 20% of the perceptual variation in 15 olfactory phenotypes, highlighting the importance of single OR genotype, ancestry, and demographic factors in variation of olfactory perception.

## Introduction

Understanding how the olfactory system detects odorants and translates their features into perceptual information is one of the fundamental questions in olfaction. Although early color vision researchers were unable to directly observe receptor responses, perceptual deficits caused by genetic variation, i.e. colorblindness, helped to show that color vision is mediated by three receptors responding to different wavelengths of light^1, 2^. Guillot, and then Amoore, extended this idea to olfaction, and proposed that cataloging specific anosmias, the inability to perceive a particular odorant, may provide similar clues linking gene and perception^3–5^. Early applications of this idea failed, presumably because olfaction relies on hundreds of receptors, and without direct observation of their responses, psychophysical tests could not untangle the fundamental rules of odor coding.

With the advent of next-generation genome sequencing to profile olfactory receptor (OR) genes and cell-based assays to identify ligands for ORs, however, receptor variation can now be matched to individuals and receptor responses can be directly observed. Humans have approximately 400 OR genes that are intact in at least part of the population, but individuals have different repertoires of pseudogenes, copy number variations (CNVs), and single nucleotide polymorphisms (SNPs) that can alter receptor responses^6–8^. While nonfunctional genes are rare in the genome (on average 100 heterozygous and 20 homozygous pseudogenes in an individual), they are significantly enriched in OR genes^9^. This provides a useful set of “natural knockouts” to examine the role of a single OR in olfactory perception—does loss of function in one OR affect odorant detection threshold, intensity, pleasantness, or character? Alternatively, the large set of ORs may redundantly encode odorant representations, such that loss of function of a single OR only rarely has perceptual consequences. Recent work suggests functional changes in a single receptor can have significant perceptual consequences, but data linking perceptual and genetic variation exist for only five ORs^10–14^.

To answer this question, we collected a larger dataset under the premise that studies of natural perceptual variation will improve our understanding of the normal translation of OR activity to odor perception. By taking advantage of high-throughput sequencing, which permits low-cost, high-coverage sequencing of the OR subgenome, we examined how perception is affected by genetic variation in the entire OR repertoire. To validate the functional link between genetic variation and perception, we used a cell-based assay to examine receptor response to associated odorants.

Here, we identified associations between genetic variation in 418 OR genes and 276 olfactory phenotypes, used cell-based functional assays to provide a mechanistic basis underlying the associations, and examined the contribution of single OR genotype, genetic ancestry, age, and sex to variation in olfactory perception.

## Results

We carried out high-throughput sequencing of the entire OR subgenome in a cohort of 332 participants previously phenotyped for their sense of smell. This dataset includes ratings of the perceived intensity and pleasantness of 68 odorants at two concentrations (**Supplementary Table 1**), detection thresholds of three odorants, and overall olfactory acuity^15^. Participants rated each stimulus twice, and the median within-subject correlation was 0.63 for intensity rating and 0.57 for pleasantness. In addition, within-participant variability was similar when ratings were collected either thirty minutes or one year apart, indicating variability is stable over time^15^.

### High-throughput sequencing of the olfactory receptor gene family

We used Illumina short-read DNA sequencing to analyze a target region consisting of 418 ORs and 256 olfactory related genes (approximately 800 kilobases), obtaining a minimum of 15x coverage for 96% of targeted bases, and identified 19,535 variants in open reading frames and contiguous regions. We validated a subset of these by comparing them to variants identified from Sanger sequencing data for 10 ORs. For eight ORs, we found greater than 95% concordance between the two sequencing methods (n ≥ 68 subjects). In contrast concordance between Sanger and high-throughput sequencing was 40% for OR10G4 and 81% for OR10G9. These ORs share a high degree of sequence similarity (OR10G4 is 96% identical to OR10G9 and 95% identical to OR10G7 at the nucleotide level according to BLAST (basic local alignment search tool)^16^), decreasing the ability to reliably map a sequencing read to a genomic location. Lower confidence in mapping (represented by lower median mapping quality (MAPQ)) associated with lower concordance between Sanger and high-throughput sequencing (Supplementary Fig. 1), potentially reflecting errors in variant calling due to read mismapping. We therefore used Sanger sequencing for OR10G4 and OR10G9. These results indicate that for ORs with high sequence similarity, alternate sequencing methods may prove necessary. However, high-throughput sequencing successfully identified sequence variants with greater than 95% accuracy for eight of the 10 ORs examined and can be expected to perform with similar accuracy on ORs where sequencing reads map with high confidence (90% of ORs have a MAPQ > 30, mapping confidence = 99.9%).

### Genetic variation in single ORs frequently associates with odor perception

We first examined the association between the genotype of single ORs and 276 different phenotypes (Fig. 1 and **Supplementary Table 2**). Eight odorant perception phenotypes (12% of the tested odors) significantly correlated with variation in a single OR locus (p < 0.05 following false discovery rate correction). Note that multiple ORs in a single locus commonly associated with a phenotype due to the structure of the olfactory subgenome (see below). These results indicate that although a given odorant typically activates multiple ORs, variation in a single OR frequently associated with perceptual features. For these top associations, OR variation was more likely to associate with the perceived intensity (88% of eight significant associations) than the perceived pleasantness of an odorant (p = 0.07 via a Binomial Test).

**Figure 1.**
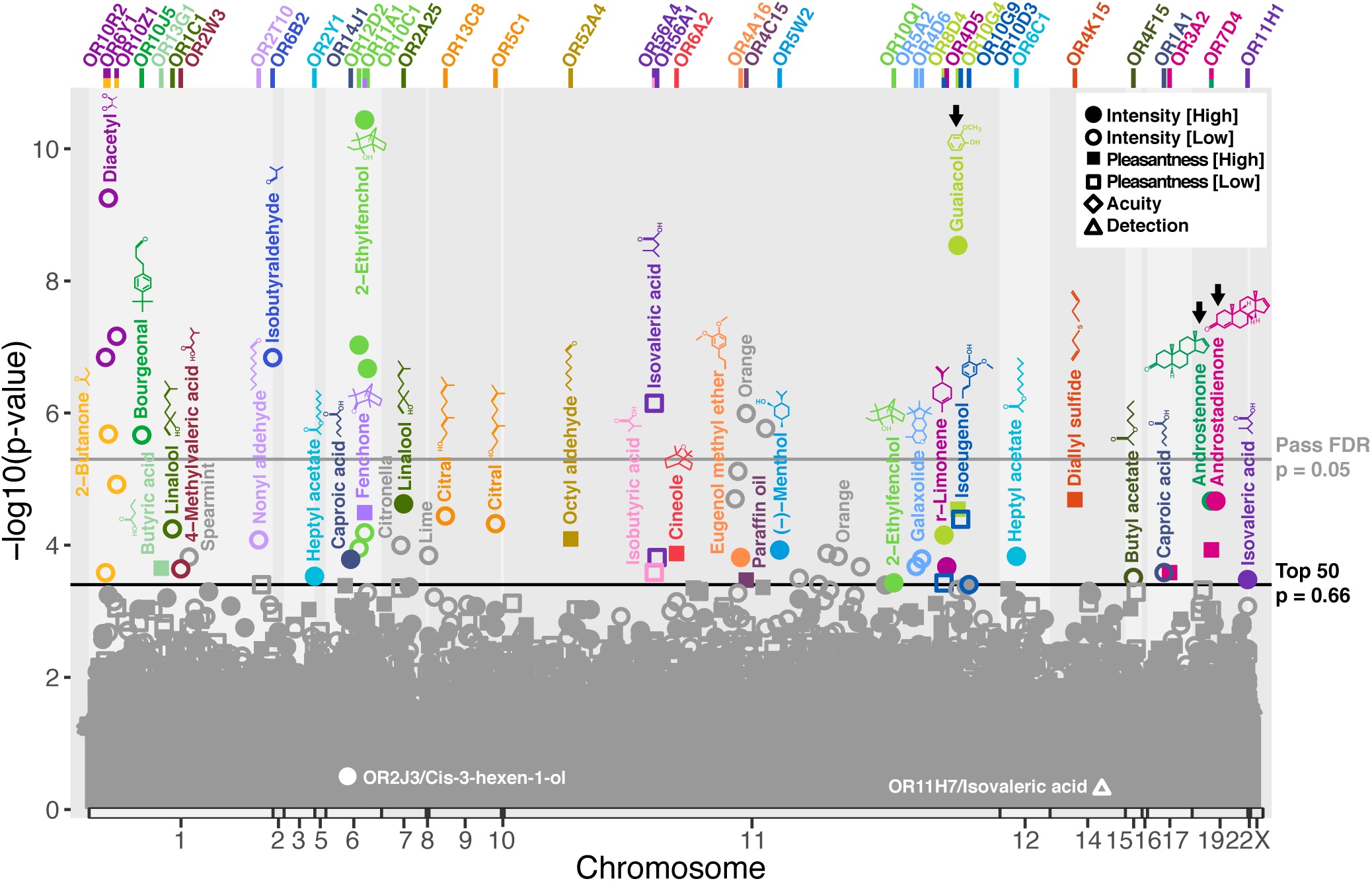
Association between OR haplotype and odorant perception. ORs are plotted by chromosomal position against association with an olfactory phenotype (-log10 p-values): perceived intensity (circles) or pleasantness (squares) rank of 68 odorants at high (closed) and low (open) concentrations, detection threshold rank of 3 odorants (triangle), or general olfactory acuity (diamond). The gray line represents p = 0.05 following multiple comparisons correction (FDR). The black line represents p = 0.66 (following FDR), the significance cutoff of the top 50 associations which were tested in cell culture (colored points). Associations with mixtures above the black line are shown in gray (spearmint, citronella, orange, and lime), because we did not analyze mixtures as ligands in our heterologous expression experiments. Points are colored by associated odorant. For loci where multiple ORs associate with an odorant, only the top association is labeled. Associated ORs are identified at the top of the graph. Previously published associations are shown in white (different dataset^10,12^) or indicated by black arrows (same dataset^11,14^).

### *In vitro* assays confirm genetic associations

Next we used a functional assay to search for a mechanistic explanation for the associations between OR genetic variation and perception. Because ORs are commonly found in clusters in the genome^17^, our association analysis had the resolution to identify regions in the genome associated with odorant perception, but could not discriminate between the causal mutation and nearby SNPs in linkage disequilibrium with the causal mutation^18–20^.

To illustrate this point, three different ORs in close proximity on chromosome 6 significantly associated with the perceived intensity of 2-ethylfenchol: OR11A1, OR12D2, and OR10C1 (Fig. 2a). The top-associated SNP (**Supplementary Table 3**), found in OR10C1, is in high linkage disequilibrium with SNPs in OR11A1 and OR12D2 (correlation > 60% in the 1000 Genomes EUR data^21,22^).

**Figure 2.**
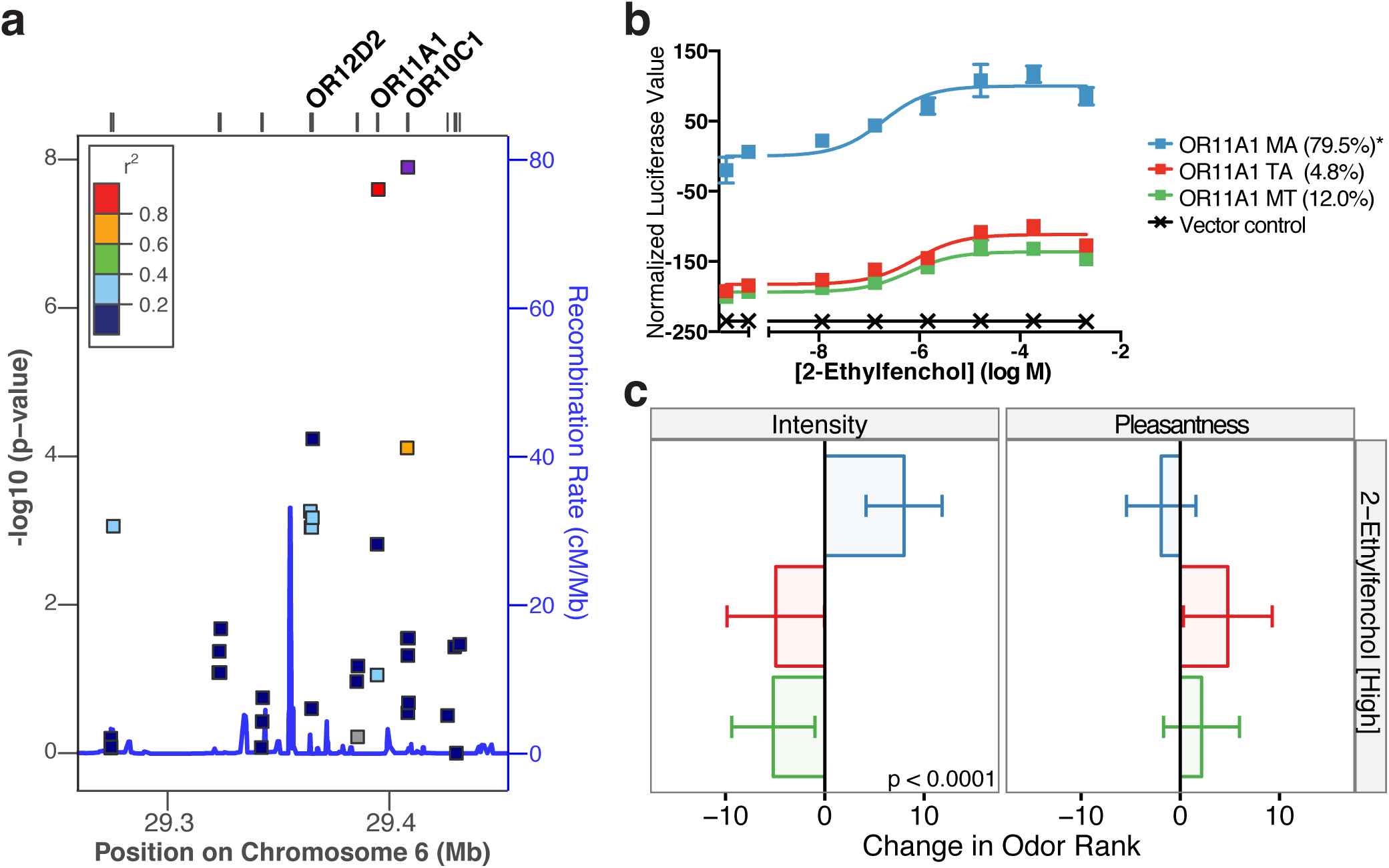
Functional variation in OR11A1 correlates with the perception of 2-ethylfenchol. (**a**) Association (-log10 p-values) between SNPs in a locus on chromosome 6 and the perceived intensity of 2-ethylfenchol. The most highly associated SNP is shown in purple, and flanking SNPs are colored according to their linkage disequilibrium with the best-associated SNP (pairwise *r*^2^ values from 1000 Genomes EUR data^21,22^). Recombination rates in this locus are shown in blue (right axis). ORs labeled at the top of the plot were tested in cell culture for their response to 2-ethylfenchol. Unlabeled lines are non-OR genes or ORs that were not tested in cell culture due to low correlation with the top SNP. (**b**) Response of OR11A1 haplotypes to increasing doses of 2-ethylfenchol. Error bars, s.e.m. of three replicates. Y-axis values are normalized to the baselined response of the reference haplotype. Letters indicate the amino acids which differ from hg19 reference haplotype (marked with an asterisk). (**c**) Changes in perceived intensity and pleasantness rank elicited by a single copy of each OR11A1 haplotype (intensity: p < 0.0001, pleasantness: p > 0.05 following FDR).

To investigate these associations, we cloned all major haplotypes in this locus with a frequency greater than 5% in our participant cohort and tested their response to increasing doses of 2-ethylfenchol using a cell-based luciferase reporter gene assay. The reference haplotype of OR11A1 (MA) was previously demonstrated to respond to 2-ethylfenchol (Fig. 2b)^14^. We found that two additional haplotypes of OR11A1, MT and TA, each with a single amino acid change from the reference, responded to 2-ethylfenchol with similar sensitivities (log(EC50 of reference MA) = −6.8, log(EC50 of MT) = −6.3, log(EC50 of TA) = −6.2). However, the MT and TA haplotypes had lower maximum responses that differed significantly from the reference MA haplotype (sum of squares test MT against MA, F(3,42) = 529.8, p < 0.0001; sum of squares TA against MA, F(3,42) = 418.9, p < 0.0001). In contrast, OR12D2 and OR10C1 did not respond to 2-ethylfenchol in the *in vitro* assay.

Furthermore, OR11A1 genotype explained 13.1% of the variance in the perceived intensity of 2-ethylfenchol (p < 0.0001 following FDR correction). Participants rated 2-ethylfenchol to be 8 ranks more intense (out of 68) for each reference MA haplotype they possessed, and 5 and 5.2 ranks less intense for each TA or MT haplotype they possessed, respectively (Fig. 2c). In addition, participants with the reference MA haplotype tended to rate 2-ethylfenchol to be less pleasant than TA or MT participants (p > 0.05 following FDR correction). These results indicate that of the three ORs found to significantly associate with the perceived intensity of 2-ethylfenchol in our analysis, OR11A1 was the best candidate for a causal receptor at this locus.

### Hyporesponsive haplotypes are associated with changes in perceived intensity or valence

We performed a similar analysis for our top 50 associated OR/odorant phenotype pairs (colored points in Fig. 1), relating the response of associated and linked ORs to odorant perception in our participants. Note that although several odor mixtures were included in the psychophysical stimulus set, we only examined associations with monomolecular odorants (gray points in Fig. 1). After examining linkage disequilibrium at each locus (Supplementary Fig. 2) and removing cases where multiple ORs from a single locus associated with the same odorant, we identified a total of 36 unique OR loci/odorant associations. Based on the false discovery rate for these top associations, we expected 66% to be false positives^23^. To discriminate false and real positive hits in our association analysis, we cloned all major haplotypes for associated ORs, in addition to at least one haplotype for any OR linked to the associated receptor (SNP correlation > 0.6, corresponding to 200 clones in total, **Supplementary Table 4**), and tested their response to the associated odorant in our heterologous assay. We tested clones that accounted for, on average, 74% of the haplotypes found in our participant population.

We found that 11 (31%) of these loci had at least one OR which responded to the associated odorant in cell culture, with a total of 14 responsive ORs (Fig. 2a, 3). Three OR/odorant pairs (OR7D4/androstenone, OR7D4/androstadienone, and OR10G4/guaiacol) were previously identified using this dataset with Sanger sequencing^11, 14^ and confirmed here via Illumina sequencing. For five of these 14 responsive ORs, OR genotype associated with perceived intensity, and subjects with less responsive haplotypes tended to rate the perceived intensity of the receptors’ ligands to be less intense than subjects with more responsive haplotypes (Fig. 2b,c, 3c,h,i, and a previously published correlation between OR7D4 function and the perceived intensity of androstenone^11^). In contrast for one OR, OR genotype associated with perceived intensity, and subjects with the more responsive haplotype rated the receptor’s ligand to be less intense (Fig. 3d). For two of these 14 ORs, OR genotype associated with perceived pleasantness, and subjects with less responsive haplotypes rated the perceived pleasantness of the associated odorant to be higher (isoeugenol) or lower (fenchone) than subjects with more responsive haplotypes (Fig. 3e,f). For three ORs, OR genotype associated with both intensity and pleasantness, and subjects with less responsive haplotypes rated the odorant as less intense and more pleasant than subjects with more functional haplotypes (Fig. 3a, and the previously published correlations between OR7D4 and OR10G4 function and the perception of androstadienone^11^ and guaiacol^14^, respectively).

**Figure 3.**
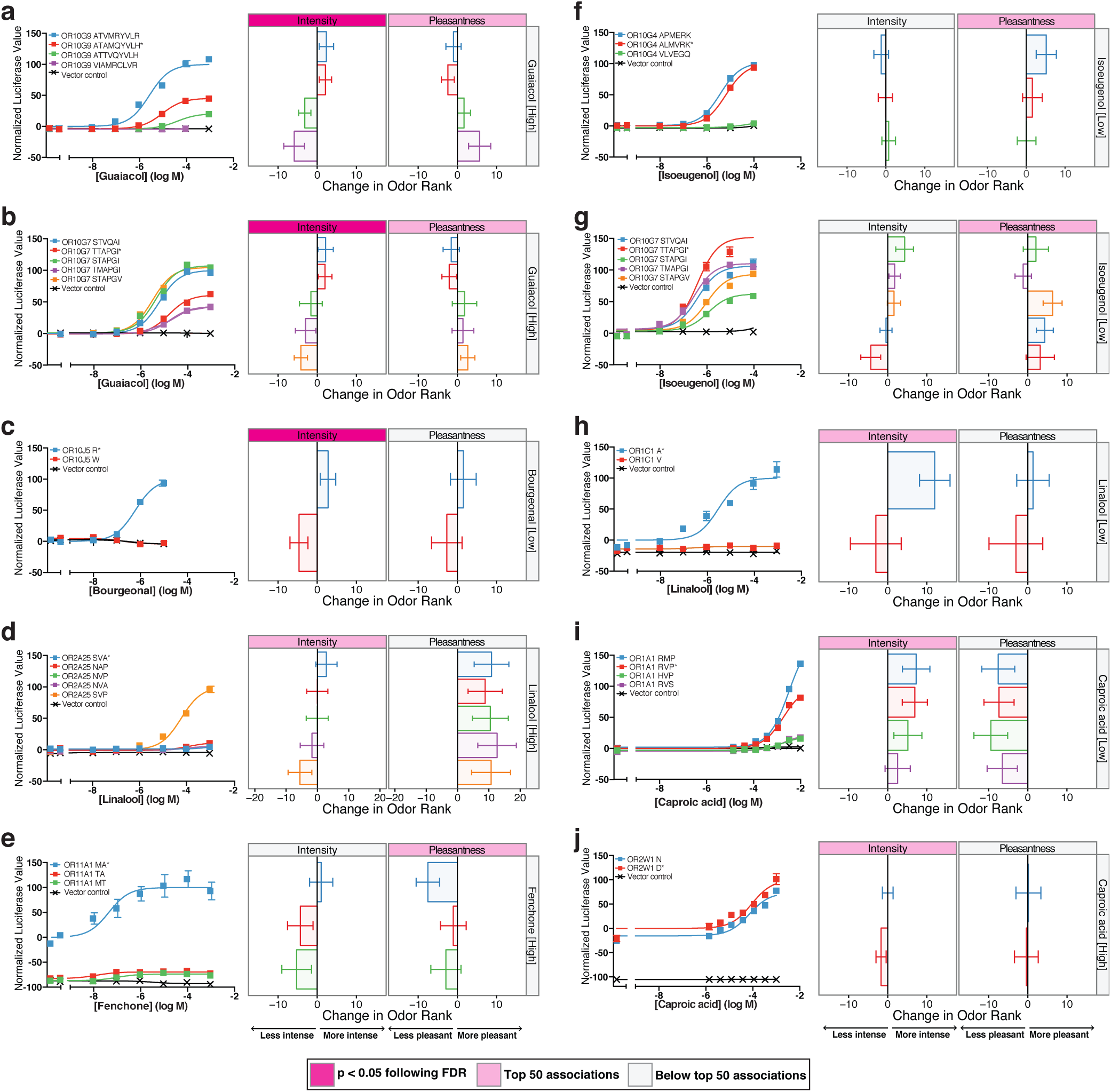
Functional variation in vitro associates with changes in perceived intensity and pleasantness. The correlation between genetic variation, OR response to odorants *in vitro*, and perceptual rankings for all OR/odorant associations with at least one responsive OR (excluding previously published OR/odorant pairs^11,14^). Left panels show the response of different OR haplotypes to increasing doses of odorant. Error bars, s.e.m. of three replicates. Y-axis values are normalized to the baselined response of either the reference or the most responsive haplotype. Right panels show the change in perceived intensity and pleasantness rank elicited by a single copy of the haplotype. Letters indicate the amino acids that differ from hg19 reference haplotype (marked with an asterisk). Pink labels indicate associations that fall within the top 50, and darker pink labels indicate associations that are significant (p < 0.05) following FDR correction.

For three ORs, haplotype function did not clearly associate with perception: similar *in vitro* function was associated with changes in perception (Fig. 3b,j) or differences in function were associated with little change in perception (Fig. 3g). These ORs may be responding to the associated odorant, while having only a small effect on perception. Overall, *in vitro* functional variation in a single OR predicted intensity or pleasantness perception, or both, for ten different OR/odorant pairs in this dataset, three of which were previously published^11, 14^.

Finally, in four cases phenotypes significantly associated with genetic variation in a single OR locus (p < 0.05 following multiple comparisons correction), but we were unable to identify an OR which responded to the associated odorant in cell culture (Fig. 4). Not all ORs have been functionally expressed in *in vitro* assay systems^24^. In order to determine if our inability to identify a causal OR was due to technical limitations, we modified the ORs in Figure 4 using conserved residues in rodent and primate orthologs as a guide and tested their response to the associated odorant. Modified OR6Y1 had nine amino acid changes from human OR6Y1 and was found in three primate species (Supplementary Fig. 3a); it responded to diacetyl, but not to 2-butanone, providing some support for the idea that OR6Y1 is the causal receptor for the association with diacetyl, although we were unable to observe its response with our assay system (Supplementary Fig. 3b).

**Figure 4.**
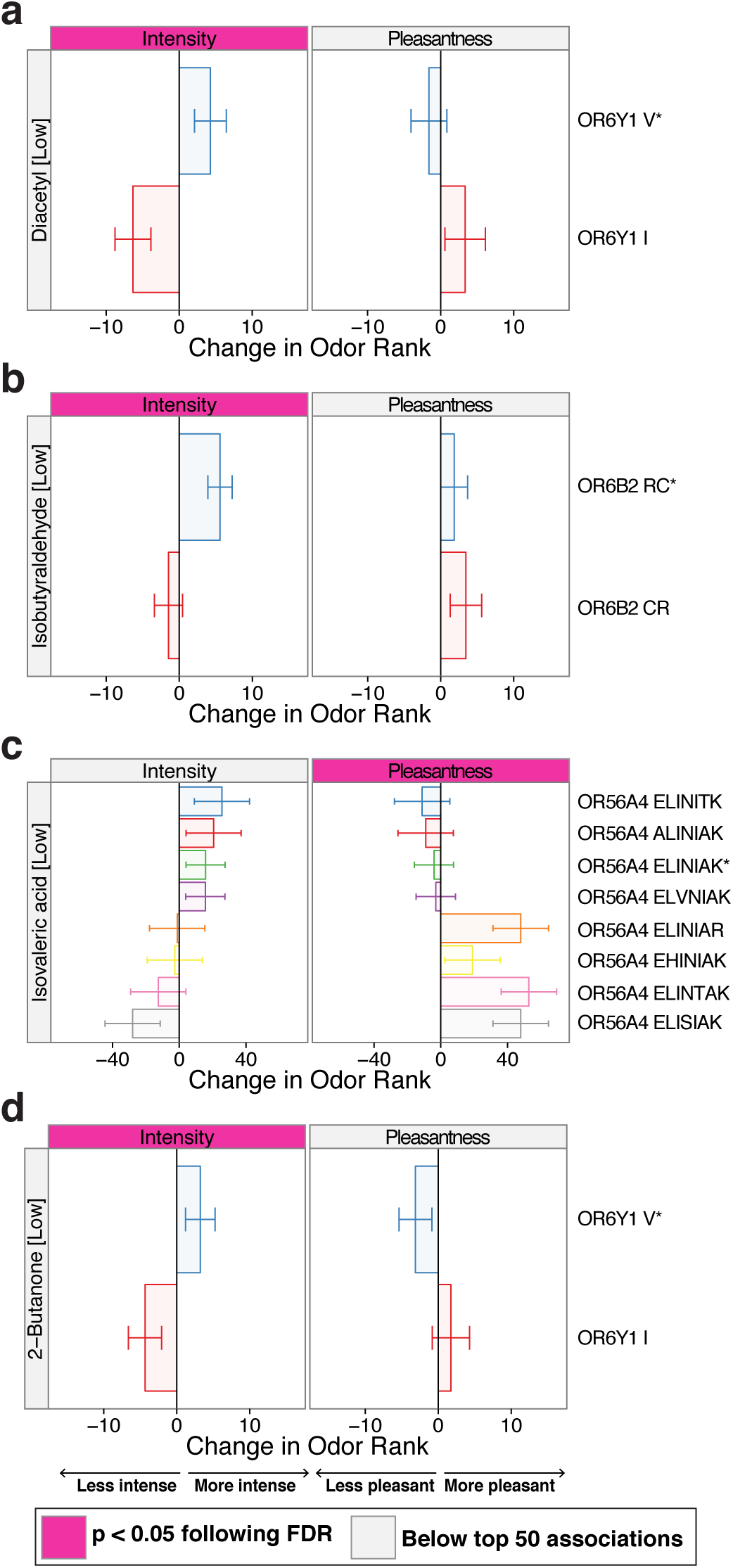
Causal ORs were not identified for all significant associations. Significant associations between genetic variation and perceptual rankings for which we were unable to identify a causal receptor. The change in perceived intensity and pleasantness rank elicited by a single copy of the haplotype is shown for OR6Y1 and diacetyl (**a**), OR6B2 and isobutyraldehyde (**b**), OR56A4 and isovaleric acid (**c**), and OR6Y1 and 2-butanone (**d**). Panels are ordered by significance. Letters indicate the amino acids that differ from hg19 reference sequence (marked with an asterisk). Pink labels indicate associations that fall within the top 50, and darker pink labels indicate associations which are significant following FDR correction.

### Genetic and demographic influences on human odor perception

We then determined how much phenotypic variance in odorant perception could be explained by various genetic and demographic factors. First, we examined the relationship between genetic ancestry and odorant perception in our participant cohort by performing a principal component (PC) analysis on all genetic data available for our subjects. Self-reported ancestry clustered when participants were plotted along the axes of the first two PCs (Supplementary Fig. 4a). This clustering was not dependent on OR genes (correlation between PC1 from all genetic data and PC1 from non-OR genetic data = 0.99, Supplementary Fig. 4b). PC1 separated Caucasians and Asians from African-Americans (self-reported), while PC2 separated Caucasians and African-Americans from Asians (self-reported).

We next examined the correlation of each PC with all 276 phenotypes. PC1 explained greater than 4% of the variance in six different phenotypes (p < 0.01 following FDR correction) (Supplementary Fig. 4c and **Supplementary Table 5**). For example, PC1 significantly correlated with the perceived pleasantness of vanillin (r = 0.28, p < 0.0001 following FDR), with self-reported Caucasians and Asians rating vanillin as more pleasant than African-Americans (Supplementary Fig. 4d). PC2 had a smaller effect on olfactory phenotypes (Supplementary Fig. 4e, p > 0.05 for all phenotypes) and explained at most 4% of the variance in spearmint perception (r = −0.20, p = 0.054 following FDR), which self-reported Asians tended to rate as more pleasant than African-Americans and Caucasians (Supplementary Fig. 4f and **Supplementary Table 5**). These results indicate that genetic ancestry is an important contributor to variation in some olfactory phenotypes.

In addition, we examined the frequency of genes that are pseudogenized in a subset of our cohort. We designated OR haplotypes with nonsense or frameshift mutations as non-functional. The median number of pseudogenized ORs in an individual was 34 out of 418 ORs that are intact in at least part of the population (Supplementary Fig. 5a), and pseudogene frequency significantly correlated with PC1, as subjects who self-reported as African-American tended to have a higher number of non-functional ORs than Caucasian and Asian subjects (r = −0.36, p < 0.0001) (Supplementary Fig. 5b). These observations are in accordance with previous work demonstrating greater pseudogene frequency in the entire genome in African versus Caucasian and Asian populations in the 1000 Genomes cohort^9^. No perceptual phenotypes significantly corre lated with OR pseudogene frequency (p > 0.05 following FDR, data not shown), indicating that number of pseudogenes did not associate with changes in detection threshold, perceived intensity, or perceived pleasantness of single odorants, or our measurement of overall olfactory acuity.

Finally, we built a model to determine how much phenotypic variance in odorant perception could be explained by four genetic and demographic factors: the genotype of the single OR that explained the most variance for each phenotype, genetic ancestry, gender, and age (**Supplementary Table 6**). The genotype of a single OR and genetic ancestry were significant contributors to several phenotypes (Fig. 5a). Age explained greater than 4% of the variance in several different phenotypes (p < 0.0001). The relative perceived intensity of nonyl aldehyde, linalool, and isobutyraldehyde decreased with age, while the relative perceived pleasantness of decyl aldehyde and the detection threshold of isovaleric acid increased with age. Note that these were changes in relative ranking, not overall decreases in sensitivity due to age. Gender explained a small percentage of variance for some olfactory phenotypes. Men tended to rate the relative perceived intensity of terpineol, banana, and citral to be higher, the perceived intensity of fir to be lower, and had a lower overall olfactory acuity than women (p < 0.002).

**Figure 5.**
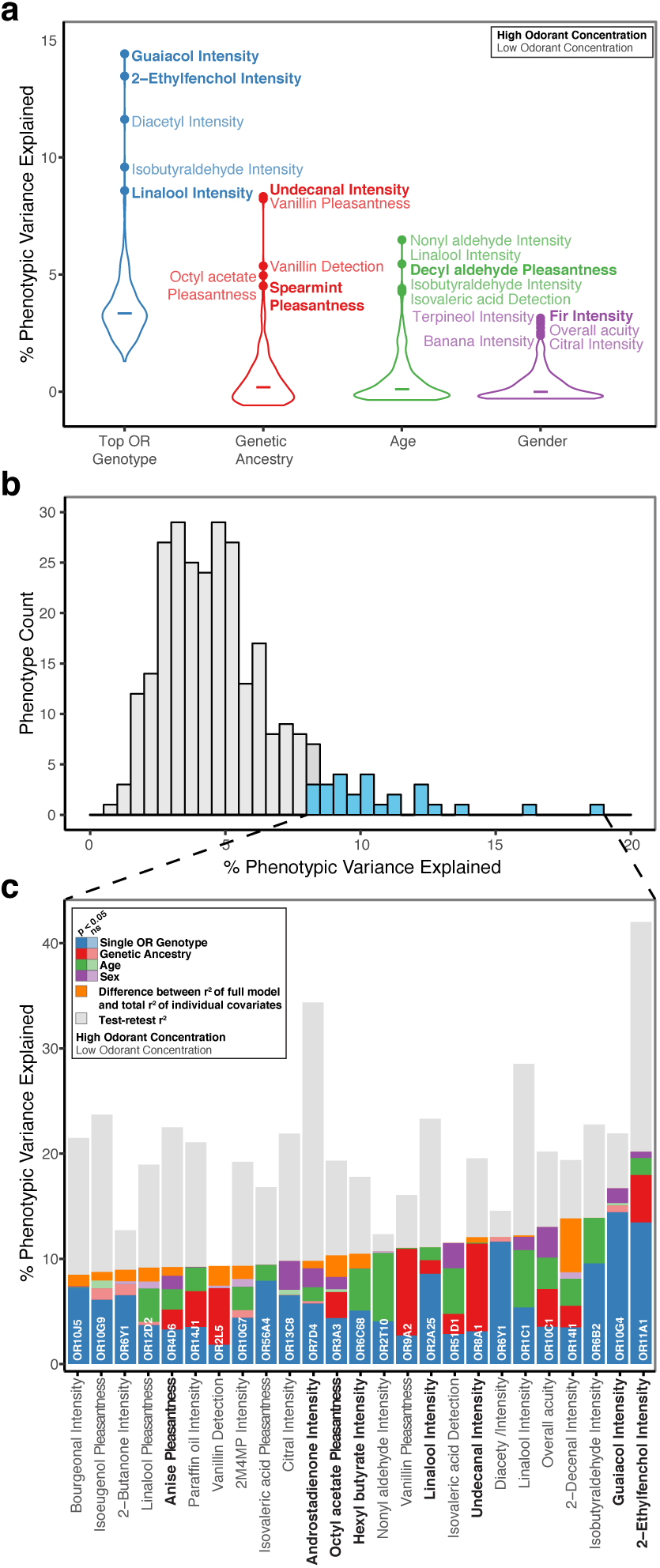
Contribution of single OR genotype, genetic ancestry, age and gender to odorant perception. (**a**) Individual contributions of single OR genotype, genetic ancestry, age, and gender to 276 odorant perception phenotypes, focusing on the OR able to explain the most variance in a particular phenotype ("Top OR Genotype"). The width of the violin plot indicates the number of phenotypes for which each factor explains that particular percentage of variance, and the line indicates the median percentage of variance explained. Top five phenotypes are labeled for each factor. (**b**) Distribution of the total percentage phenotypic variance explained by the full model. Bars in blue illustrate the top 25 perceptual phenotypes (p < 0.0001 following FDR), and the relative contribution of all 4 genetic and demographic factors to variance in these phenotypes is shown in (**c**). Covariates that significantly alter the linear model are shown in full color (p < 0.05), as determined by a one-way analysis of variance comparing the complete model (with all four covariates) to a model excluding the covariate. Gray bars indicate the total explainable variance for each phenotype, as determined by its test-retest value. Bold labels indicate high odorant concentration, and plain labels indicate low odorant concentration.

The median percentage of total phenotypic variance we could explain using single OR genotype, genetic ancestry, gender, and age was 4.77% (Fig. 5b). We then assessed the relative contribution of all four factors to the 25 phenotypes for which we could explain the most variance (p < 0.0001) (blue in Fig. 5b and shown in more detail in Fig. 5c). The test-retest correlation for a phenotype provided an upper bound for the amount of perceptual variance we could explain (shown in gray Fig. 5c)^25^. Note we did not measure test-retest correlation values for detection thresholds, as these measurements were only collected once. For some perceptual phenotypes our model accounted for over 80% of the explainable variance, such as the perceived intensity of diacetyl, where single OR genotype is the main contributor, and the perceived intensity of nonyl aldehyde, where age is the main explanatory variable. For other phenotypes, such as the perceived intensity of 2-ethylfenchol and linalool, OR genotype, genetic ancestry, age, and gender accounted for less than half of the explainable variance. These results indicate that while genetic variation in a single OR is an important contributor to perceptual variance, methods that consider multiple ORs may allow us to account for more of the explainable variance in a particular phenotype.

## Discussion

We performed a large-scale examination of the relationship among OR genetic variation, OR activation, and odorant perception. We examined 36 different OR loci/odorant pairs in cell culture, demonstrated that at least one OR from 11 of these loci responded to the associated odorant, and found that OR response *in vitro* matched the perceived intensity or pleasantness of the associated odorant for 10 different OR/odorant pairs. As three of these pairs have been previously published^11, 14^, here we describe seven new cases where genetic variation in a single OR predicts intensity or pleasantness perception. To our knowledge, this is the first large-scale examination of the association between the entire OR subgenome and a large group of olfactory phenotypes.

Our results have important implications for olfactory coding. First, we demonstrated that the perception of 13% of the 68 odors tested were tied to genetic and functional variation in a single OR. This increases the known cases directly linking OR activation to odorant perception from five to 12^10–14^ and places a constraint on the amount of redundancy in the combinatorial code. These cases provide valuable tools to explore and manipulate the olfactory code using agonists or antagonists and to examine the contribution of the activation of individual ORs to the coding of odorant information.

Second, loss of function of an OR correlated with a decrease in the perceived intensity of the ligand in eight out of ten cases in this dataset, in accordance with previous studies^11, 13, 14^. The specificity of the effect is inconsistent with a proposed model where the bulk of ORs encode odorant identity and a subset of broadly tuned receptors encode odorant intensity^26^. Another proposed model of intensity coding is that the total number of ORs activated at a given concentration encodes odorant intensity. While the number of functional ORs correlated with the relative intensity within one odorant, this may not apply across odorants as several odorants that activate a large number of OR types *in vitro* are not particularly intense (eugenol), while odorants that appeared to activate only a few receptors are particularly potent (thiols). However, these observations may be explained by difficulties in comparing concentrations across *in vitro* and *in vivo* studies. The repertoire of activated ORs changes with odorant concentration, and therefore genetic variation in an OR will be perceptually relevant only at certain concentrations. We tested two concentrations of each odorant, and found that perceived intensity was associated with genetic variation in a particular OR more strongly for one concentration. This is consistent with work showing that different receptors in *Drosophila melanogaster* are necessary for perception in different concentration ranges^27^.

Third, loss of function in an OR correlated with the perceived pleasantness of the ligand in four out of ten cases. Unlike the correlation with perceived intensity, loss of function led to reduced pleasantness in one case (isoeugenol) and increased pleasantness in three cases (fenchone, guaiacol, and androstadienone). For most associations (nine out of ten), OR variation correlated with changes in both intensity and pleasantness, although the effect on the latter was usually smaller (below the top 50 associations). Rarely (one case out of ten) was variation in OR function related to pleasantness changes with no associated changes in perceived intensity (isoeugenol in Fig. 3f). One possibility is that the change in intensity is driving pleasantness, but without knowing how pleasantness changes across a full range of concentrations, we cannot test this hypothesis here.

Finally, we found that genetic variation in a single receptor had a larger effect on intensity and pleasantness than on detection threshold. Previous studies focused on the correlation between OR genetic variation and differences in detection threshold^10, 12, 28,^ but in our dataset, no single OR associated with the detection threshold of vanillin, isovaleric acid, or pentadecalactone at a significance level that warranted testing in cell culture. Poor genotyping frequency prevented us from testing a published association between OR11H7 and isovaleric acid detection threshold^10^ (Fig. 1). However, the genotype of OR56A4 significantly associated with the perceived pleasantness of isovaleric acid, but not its detection threshold. Similarly, OR7D4 genotype explains more variance in the perceived intensity of androstenone than its detection threshold^11^. Differences in phenotype measurements may also account for our failure to find a previously identified association between OR2J3 genotype and cis-3-hexenol perception, as the original study examined detection threshold and here we measured perceived intensity^12^ (Fig. 1).

The high linkage disequilibrium that characterizes many OR loci makes identifying causal ORs difficult using association analyses of the scale discussed here^17, 29^. High resolution analyses capable of dissecting these loci would require perceptual data from an extremely large number of subjects. For example, genome-wide association studies (GWAS) examining asparagus urine odor anosmia and cilantro preference in 4,742 and 26,455 subjects, respectively, were still unable to narrow down causal regions beyond a cluster of ORs^30, 31^. To overcome these limitations, we used a cell-based assay to identify the response of the associated and linked ORs to odorants.

Heterologous assays have identified ligands for only 12% of the roughly 400 intact human ORs to date^32^, suggesting that these assays need to be improved to functionally express all human ORs. Based on this success rate, the expected rate of identifying a causal OR underlying an association was around 12%. Because the false discovery rate for our top 50 associations was 66%, among the 36 OR/odorant pairs examined, we expected that 12 would be true positives and 24 would be false positives. We in fact found 11 cases where the associated OR/odorant pair was active *in vitro*, which was much higher than the expected 12% (approximately one case) given previous work in heterologous assays. These results suggest that the receptors that are relevant to perception are enriched in the set of receptors that respond in the *in vitro* assay. ORs which are functional *in vitro* are also more likely to be found in the set of receptors expressed in the human olfactory epithelium (OE)^33^. Furthermore, deorphanized (as listed in^33^) or perceptually relevant (^10–14^, Fig. 2, 3) ORs are expressed at levels 1.5 times higher than other intact ORs^34^ (p < 0.0001 for both via a binomial test). One possibility is that a large fraction of intact human ORs are nonfunctional *in vivo* and are not expressed in the OE. In summary, our results indicate that cell-based assays are a useful proxy for identifying behaviorally relevant ORs that are expressed in the OE and whose activation can be directly tied to perception. Despite these successes, there are certainly cases where *in vitro* results do not predict perception—*in vitro* assays lack critical components of the OE, including proteins in the mucous layer which transport and modify odorant molecules^24^. Cell-based assays were unable to identify a responsive OR for some loci for which we have strong association data (Fig. 4a), potentially due to either the association being spurious or the receptor failing to function in the cell-based assay. In one instance, a modified version of OR6Y1 derived from rodent and primate orthologs responded to diacetyl, supporting the idea that, although the human receptor did not function properly in the *in vitro* assay (Supplementary Fig. 3), it was the causal receptor underlying this association. Examining cases where the *in vitro* assay does not match perceptual outcomes may provide guidance on how to improve these assays in the future.

We found that several olfactory phenotypes were significantly influenced by ancestry. Ancestry is a common confounding factor in association studies that incorporate different subpopulations^35^. Although most previous studies examining the association between OR genotype and odorant perception circumvented this problem by using a homogenous participant cohort^10, 28, 36,^ studies conducted in diverse populations indicate that ancestry may be a significant factor in odorant perception^29^. A previous examination of the relationship between participant demographics and odorant perception in our cohort demonstrated that self-reported ancestry (African-American, Caucasian, Asian) significantly correlated with some perceptual phenotypes^15^. Here, we extended these results by quantifying ancestry using all genetic data available for our subjects (roughly 19,000 SNPs in both ORs and 256 additional genes), bypassing self-reporting and allowing us to incorporate subjects who self-reported their ancestry as “Other”. Although we found no significant associations between SNPs in the 256 non-OR genes and the tested phenotypes (**Supplementary Table 3**), the SNPs were useful in establishing ancestry. We found that ancestry was able to explain a significant portion of the variance in undecanal and 2-decenal and the perceived pleasantness and detection threshold of vanillin (also shown in^15^), among others. From these data alone, we were unable to determine if these differences in odorant perception were due to unknown genetic, cultural, or social factors that co-segregate with ancestry. In conjunction with our finding that OR pseudogene frequency was higher in subjects who self-identified as African-American, this work demonstrates the importance of considering ancestry when studying odorant perception in diverse populations. Models incorporating OR genotype, ancestry, age, and gender accounted for over 70% of the explainable variance (test-retest correlation) for some olfactory phenotypes (guaiacol, diacetyl, and nonyl aldehyde) and less than half of the explainable variance for others (2-ethylfenchol, linalool, and androstadienone). These results indicate that considering the contribution of multiple ORs may be useful in explaining more of the variance for some olfactory phenotypes. While the contribution of genetic and demographic factors to variation in odorant perception has been previously examined, this is the first investigation to examine the entire OR subgenome, quantify ancestry using genetic variation, and relate these and other demographic factors to a large group of olfactory phenotypes. This work also highlights the necessity of considering the contribution of multiple ORs to odorant perception.

Although we know the OR gene family is characterized by a large amount of genetic and functional variation, the combinatorial nature of the olfactory code and our limited knowledge of OR/odorant pairs makes it difficult to translate this variation to differences in perception. Here we focused on the perceptual consequences of loss-of-function of individual ORs, demonstrating that intensity and pleasantness coding for some odors is not redundant and that loss of function in a receptor reduces the perceived intensity of the receptor’s ligand. Similar studies on colorblindness, a condition in which genetic variation alters color perception, helped determine the tuning of the three photoreceptors to different wavelengths of light. Deciphering the quantitative representation of color by photoreceptors allowed us to digitize color information so that it can be sent and stored without degradation, as well as develop representations of color space that outline how wavelengths of light can be combined to make novel colors. Understanding how the olfactory receptors encode odors should lead to similar advances in olfaction: namely digitizing odors and identifying agonists or antagonists of receptors that can produce any desired olfactory percept from a small set of primary odors.

## Methods

### Psychophysical testing

Collection of psychophysical data was previously reported by Keller et al. and approved by the Rockefeller University Institutional Review Board^11, 15^. Briefly, 391 subjects rated both the intensity and pleasantness of 66 odors at two concentrations (designated “high” and “low”) and two solvents on a scale of 1 to 7, 1 being “extremely weak” or “extremely unpleasant” and 7 being “very strong” or “extremely pleasant”. The high and low odorant concentrations were intensity-matched to 1/1,000 and 1/10,000 dilutions of 1-butanol, respectively. For six odorants, pure odorant was rated as less intense than the 1/1000 dilution of 1-butanone, and these were not diluted for testing (odors described in **Supplementary Table 1**). In addition, the detection thresholds for three odorants (pentadecalactone, vanillin, and isovaleric acid) were determined for each subject.

Subjects rated the intensity and pleasantness of each odorant/concentration twice. Within-subject variability in odorant rating was determined by calculating the correlation between the first and second rating of all odorants. Test-retest correlation was calculated by examining the correlation between the first and second rating for each olfactory phenotype where duplicate trials were run. For each subject, the average intensity and pleasantness ratings at each odorant concentration (low and high) were ranked from 1 to 68, such that the odorant with the highest rated intensity for a concentration was ranked as 68 and the odorant with the lowest rated intensity was ranked as 1. Solvents (propylene glycol and paraffin oil) were rated three times at a single concentration, which was averaged and included in the ranking with the other 66 odorants at both concentrations. Ranking on a per-subject basis controlled for the different use of the rating scale among subjects. Detection thresholds were ranked on a per-odorant basis, such that the subject with the highest detection threshold for a particular odorant received a ranking of 1 and the subject with the lowest detection threshold for an odorant received a ranking of 391. Three measurements were used to calculate general olfactory acuity: percentage of odorants where the high concentration was rated as more intense than the low concentration, percentage of odorants where the high concentration was rated as more intense than solvent, and percentage of odorants where the low concentration was rated as more intense than the solvent. All three measurements were ranked on a per-task basis, the ranks were averaged for each subject, and finally this average was expressed as a rank among all subjects from 1 (lowest acuity) to 391 (highest acuity). Therefore, 276 different phenotypes were examined: perceived intensity and pleasantness rank of 66 odorants at two concentrations, two solvents at one concentration but included in the ranking for both high and low odorant concentration, detection threshold rank of three odorants, and overall olfactory acuity.

### Sequencing sample preparation and genotyping

Genomic DNA was prepared from venous blood samples with the PAXgene Blood DNA kit (Qiagen). DNA was sheared (Covaris) and ligated to adapters necessary for both sequencing and barcoding samples using the TruSeq kit (Illumina). The OR subgenome was captured using an Agilent SureSelect Target Enrichment kit custom-designed to enrich for the open reading frame of ORs and human orthologs of 256 additional genes found to be highly expressed in mouse olfactory sensory neurons (unpublished data) (SureSelect ELID 0352781, **Supplementary Table 7**)^37^. Paired-end sequencing was carried out on 332 participants using an Illumina GAIIx with a read length of 2x75 basepairs. Each sample was individually barcoded and twelve samples were multiplexed per lane.

Sequence variants were identified using a custom-made pipeline that followed the current best practices recommended for variant detection by the Broad Institute^38, 39^. Reads were aligned to the hg19 reference genome using BWA^40^, and the Genome Analysis Toolkit (GATK)^41^ was used to remove PCR duplicates, realign reads around insertions and deletions (indels), recalibrate base quality scores, and genotype variant sites (SNPs and indels) across all 332 subjects simultaneously using variant quality score recalibration (SNPs) or standard hard filtering (indels). SNPs were phased with SHAPEIT^42^ (excluding SNPs genotyped at a frequency < 95% and SNPs which deviated from Hardy-Weinberg equilibrium (p < 0.00001))^20^, and a custom-written R^43^ script was used to translate the phased variant call file into 836 full-length haplotypes (418 ORs x a maternal and a paternal haplotype) for each subject. Finally, Sanger sequencing was used to confirm the sequence of 10 ORs in at least 68 subjects. Genomic DNA was amplified with Phusion DNA Polymerase (Thermo Fisher Scientific) or EmeraldAmp PCR Master Mix (Clontech) using primers up- and downstream of the OR’s open reading frame (**Supplementary Table 8**). PCR products were purified by ultrafiltration on a vacuum manifold (NucleoFast 96 PCR, Machery Nagel) and sequenced (ABI 3730XL) at the University of Pennsylvania DNA Sequencing Facility.

Putative OR pseudogenes were identified by annotating variants using Variant Effect Predictor to determine type and impact of mutations^44^. The sum of OR haplotypes with nonsense or frameshift mutations was determined for each individual.

### Association Analysis

The association between olfactory phenotypes and single OR genotype was analyzed using multiple linear regression to regress the haplotype count (0, 1, or 2) of individual ORs against all 276 phenotype measurements using the R statistical package. The analysis was limited to haplotypes found at a frequency greater than 5% in our cohort and ORs with low genotype frequency were removed (29 ORs: OR4F5, OR2T8, OR2T7, OR2T5, OR2T29, OR2T35, OR2T27, OR5AC1, OR5H8, OR2A4, OR10AC1, OR2A42, OR2A7, OR2A1, OR4F21, OR51A2, OR52Z1, OR52N1, OR5G3, OR8G1, OR11H12, OR11H2, OR4Q2, OR11H7. OR4E1, OR4F4, OR1D5, OR1D4, OR4F17). Note the ORs eliminated did not have any single SNPs which significantly associated with odorant perception (see below for details on SNP association analysis). To correct for population structure (ancestry), the first two PCs calculated from all genetic data were incorporated as covariates in the linear model^35^. Principal components were calculated using the SNPRelate^45^ package in R (after removing SNPs genotyped at a rate < 95% and SNPs in linkage disequilibrium > 0.5)^20^. P-values were corrected for multiple comparisons using false discovery rate^23^. Although our ranked data is not normally distributed, linear regression was used to find the coefficients for each haplotype. Genotype/phenotype association was also analyzed using a Kruskal-Wallis test, which demonstrated that p-values between the parametric and non-parametric test were significantly correlated (*r*^2^ = 0.77, p < 0.0001).

To analyze phenotype association with the genotype of single SNPs, individual SNP counts were regressed against phenotype measurements, and the first 2 PCs were incorporated as covariates to correct for ancestry in the study population using PLINK^46, 47^. The analysis was limited to variants with minor allele frequencies greater than 5% that did not significantly deviate from Hardy-Weinberg equilibrium (p > 0.00001)^20^. P-values were corrected for multiple comparisons using false discovery rate. To examine linkage disequilibrium among SNPs in OR loci, SNP genotype correlations were calculated from 1000 Genomes data (European population) using LocusZoom^21, 22^.

### Contribution of ancestry, age, and gender to perception

First, the correlation between olfactory phenotypes and the first two PCs of genetic variation was calculated, and p-values were corrected for multiple comparisons using false discovery rate. The total contribution of single OR genotype, ancestry, age, and gender to phenotype variability was calculated by regressing the count of individual OR haplotypes against all 276 phenotypes and incorporating the first two PCs calculated from all genetic data, age (in years), and gender as covariates (full model). P-values were corrected for multiple comparisons using false discovery rate. To calculate the individual contribution (*r*^2^) of single OR genotype, ancestry, age, and gender to each phenotype, the percentage of variance explained by a linear model in which each covariate was removed was compared to the full linear model. To determine if a covariate significantly altered the linear model, a one-way analysis of variance was used to compare models with and without the covariate.

### Consensus odorant receptor

The online version of MAFFT version 7 was utilized to create OR6Y1, OR6B2, and OR56A4 consensus proteins based on the alignment of the orthologs found in *Homo sapiens, Gorilla gorilla gorilla, Pan paniscus, Pan troglodytes, Pongo abelii, Macaca mulatta, Mandrillus leucophaeus, Callithrix jacchus, Microcebus murinus, Rattus norvegicus*, and *Mus musculus*^48, 49^. The consensus genes were then designed for expression in human cells using the IDT Codon Optimization Tool and synthesized as a standard IDT gBlocks Gene Fragment.

### OR cloning

OR haplotyes for functional testing in cell culture were generated by cloning the respective sequence from pooled genomic DNA, from a specific subject, or by generating polymorphic SNPs by site-directed mutagenesis using overlap extension PCR^50^. The open reading frame of each OR was amplified with Phusion polymerase and cloned into the pCI vector (Promega) containing the first 20 amino acids of human rhodopsin^51^. Cloned sequences were confirmed by Sanger sequencing (ABI 3730XL) at the University of Pennsylvania DNA Sequencing Facility.

### Luciferase assay

The Dual-Glo Luceriferase Assay System (Promega) was used to measure *in vitro* OR activity as described previously^52, 53^. Hana3A cells were co-transfected with OR, a short from of receptor transporter protein 1 (RTP1S)^54^, the type 2 muscarinic acetylcholine receptor (M3-R)^55^, Renilla luciferase driven by an SV40 promoter, and firefly luciferase driven by a cyclic AMP response element. 18-24 hours post-transfection, ORs were treated with medium or serial dilutions of odorants spanning 1nM to 1mM in triplicate. Four hours after odorant stimulation, luciferase activity was measured using the Synergy 2 (BioTek). Normalized luciferase activity was calculated by dividing firefly luciferase values by Renilla luciferase values for each well. Results represent mean response (for 3 wells) +/-s.e.m. Responses were fit to a three-parameter sigmoidal curve. An odorant was considered an agonist if the standard error of the logEC50 was less than 1 log unit, the 95% confidence intervals for the top and bottom parameters of the curve did not overlap, and the extra sum-of-squares test confirmed that the odorant activated OR-transfected cells more than empty-vector-transfected cells. An extra sum-of-squares was also used to determine if one model fit the data from two haplotypes better than two separate models. Data were analyzed with GraphPad Prism 6.

## Acknowledgements

This work was supported by R03 DC011373 and R01 DC013339 to J.D.M; T32 DC000014 and F32 DC014202 to C.T.; DC005782, DC012095 and DC014423 to H.M.; and 2017/00726-2 to M.N. from the São Paulo Research Foundation (FAPESP). A portion of the work was performed using the Monell Chemosensory Receptor Signaling Core and Genotyping and DNA/RNA Analysis Core, which were supported, in part, by funding from the US National Institutes of Health NIDCD Core Grant P30 DC011735. Collection of psychophysical data was supported by UL1 TR000043 from the Clinical and Translational Science Award program at the National Center for Advancing Translational Sciences. L.B.V. is an investigator of the Howard Hughes Medical Institute.

## Author Contributions

J.D.M., A.K., H.M., and L.B.V. conceived and designed the project. C.T., N.R.M., L.L.S., J.R.W., M.N., N.K., and J.D.M. performed research. A.K. collected the psychophysical data and provided DNA samples under the supervision of L.B.V. C.T. carried out the analysis and wrote the paper with input from all authors. J.D.M. supervised the project.

---

**Supplementary Figure 1.**
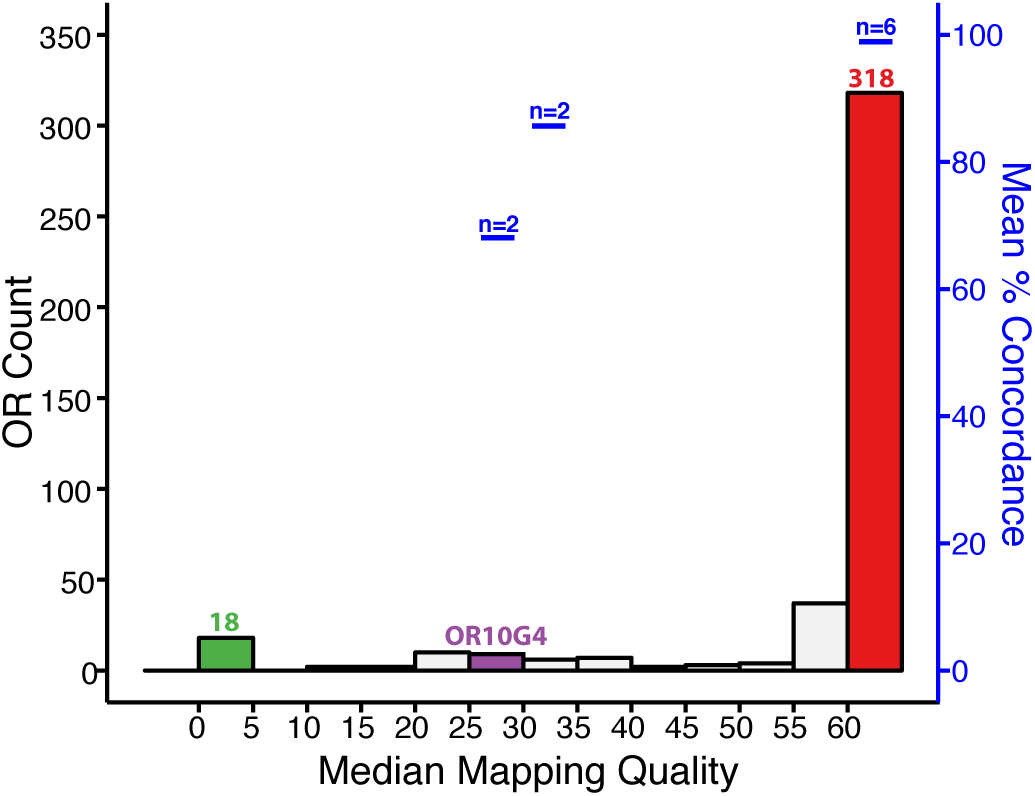
Median mapping quality for each OR. Distribution of the median mapping quality (MAPQ) calculated for all reads for 100 subjects (left axis), and the average percent concordance between Illumina and Sanger sequencing for 10 ORs of various mapping qualities (right axis).

**Supplementary Figure 2.**
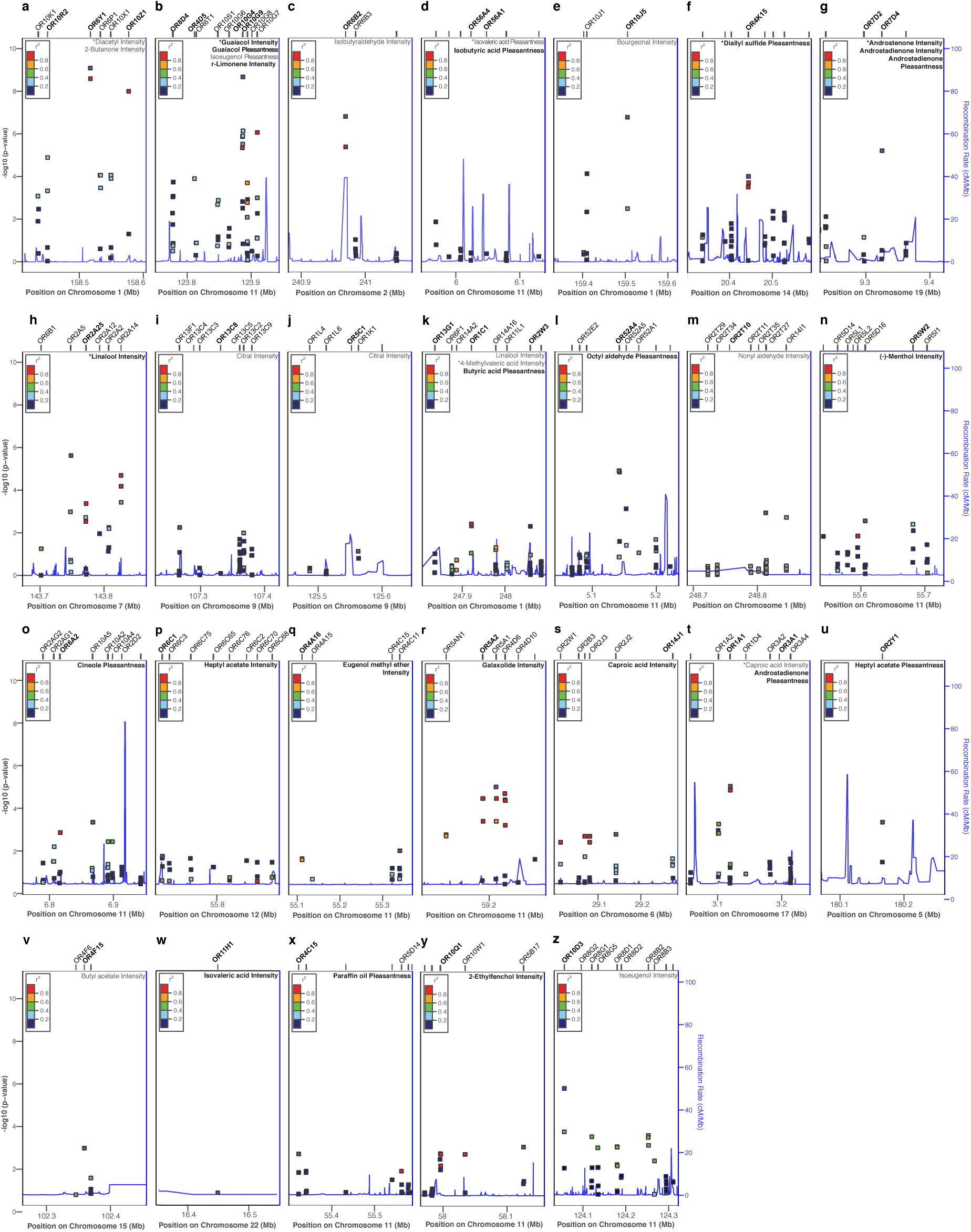
Genetic variation in OR clusters associates with odorant perception. Association (-log10 p-values) between SNPs in 27 different OR loci and the 38 perceptual phenotypes shown in the top right corner of each panel (note one locus associated with two phenotypes is shown in Fig. 2a). The most highly associated SNP is shown in purple, and flanking SNPs are colored according to their linkage disequilibrium with the best-associated SNP (pairwise *r*^2^ values from 1000 Genomes EUR data^21,22^). Recombination rates in each locus are shown in blue (right axis). ORs labeled at the top of the plot were tested in cell culture for their response to the associated odorant, and the associated ORs are shown in bold. Unlabeled lines are non-OR genes or ORs that were not tested in cell culture due to low correlation with the top SNP.

**Supplementary Figure 3.**
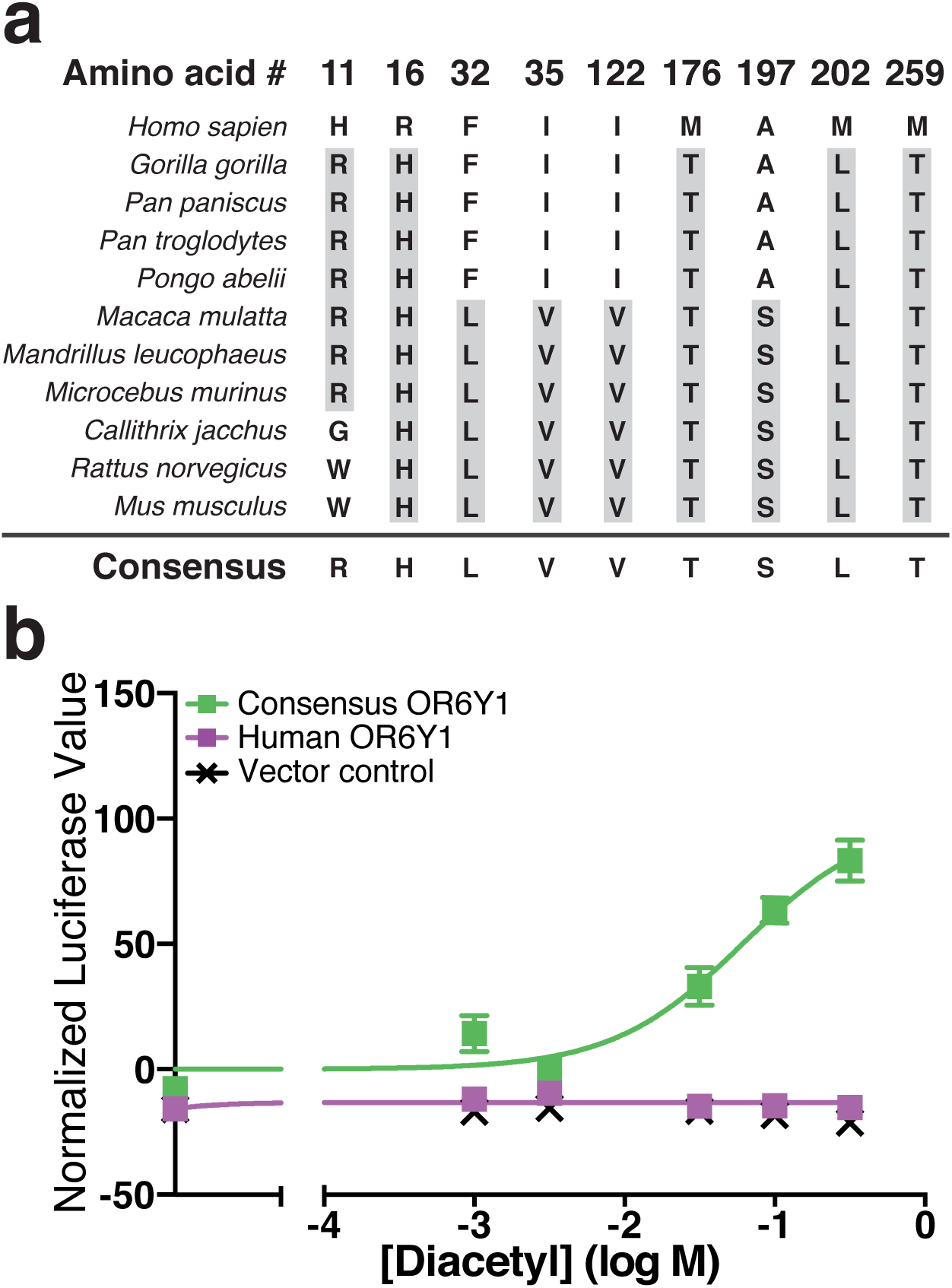
Modified OR6Y1 responds to diacetyl. (**a**) Comparison of human OR6Y1 to orthologous receptors from 10 species. The most common amino acid for each position, highlighted in gray, was used to make a consensus receptor with 9 amino acid changes from human OR6Y1. (**b**) Dose-response curves for human and modified OR6Y1. Responses are from cells transfected with either a plasmid encoding the indicated odorant receptor or an empty vector stimulated with diacetyl. Error bars, s.e.m. of three replicates.

**Supplementary Figure 4.**
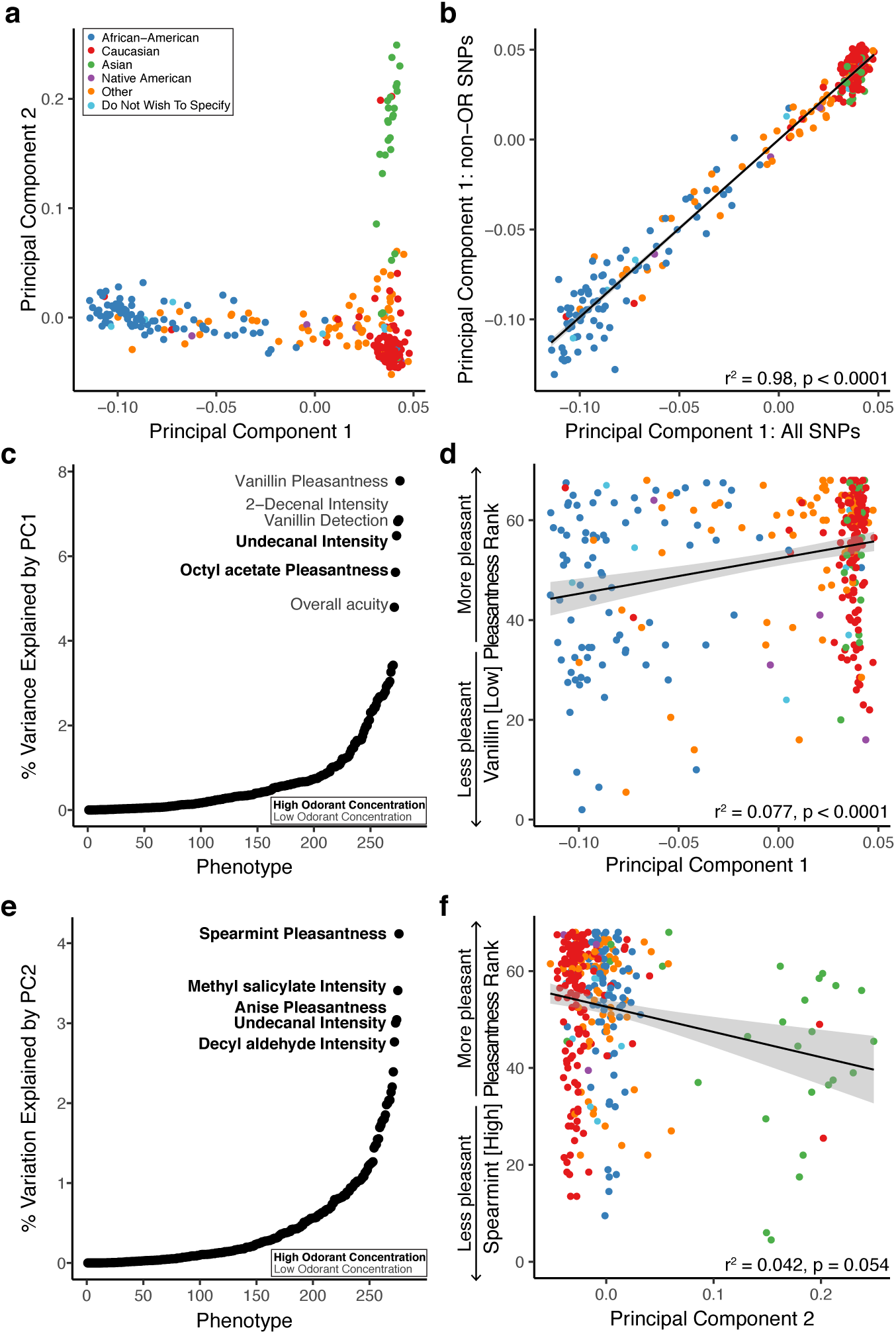
Genetic ancestry correlates with olfactory perception. (**a**) Self-reported ancestry clustered when participants were plotted according to their eigenvectors for the first two principal components (PCs) calculated from all available genotype data for our subject cohort. (**b**) Correlation between the first PC of genetic variation calculated using SNPs from all targeted genes (OR and non-OR genes) and the first PC calculated using SNPs identified in non-OR genes only (*r*^2^ = 0.98, p < 0.0001). (**c**) Percent variance explained by PC1 for all 276 phenotypes (ordered by percent variance explained). PC1 explains greater than 4% of the variance for 6 phenotypes (labeled) (p < 0.01 following FDR). Bold labels indicate high odorant concentration, and plain labels indicate low odorant concentration. (**d**) Correlation between PC1 and the perceptual ranking for the pleasantness of vanillin (*r*^2^ = 0.077, p = 0.0001). (**e**) Percent variance explained by PC2 for all 276 phenotypes (ordered by percent variance explained). The top 5 phenotypes are labeled (*r*^2^ > 0.027, p > 0.05 following FDR). Bold labels indicate high odorant concentration, and plain labels indicate low odorant concentration. (**f**) Correlation between PC2 and the perceived pleasantness of spearmint (*r*^2^ = 0.042, p = 0.054).

**Supplementary Figure 5.**
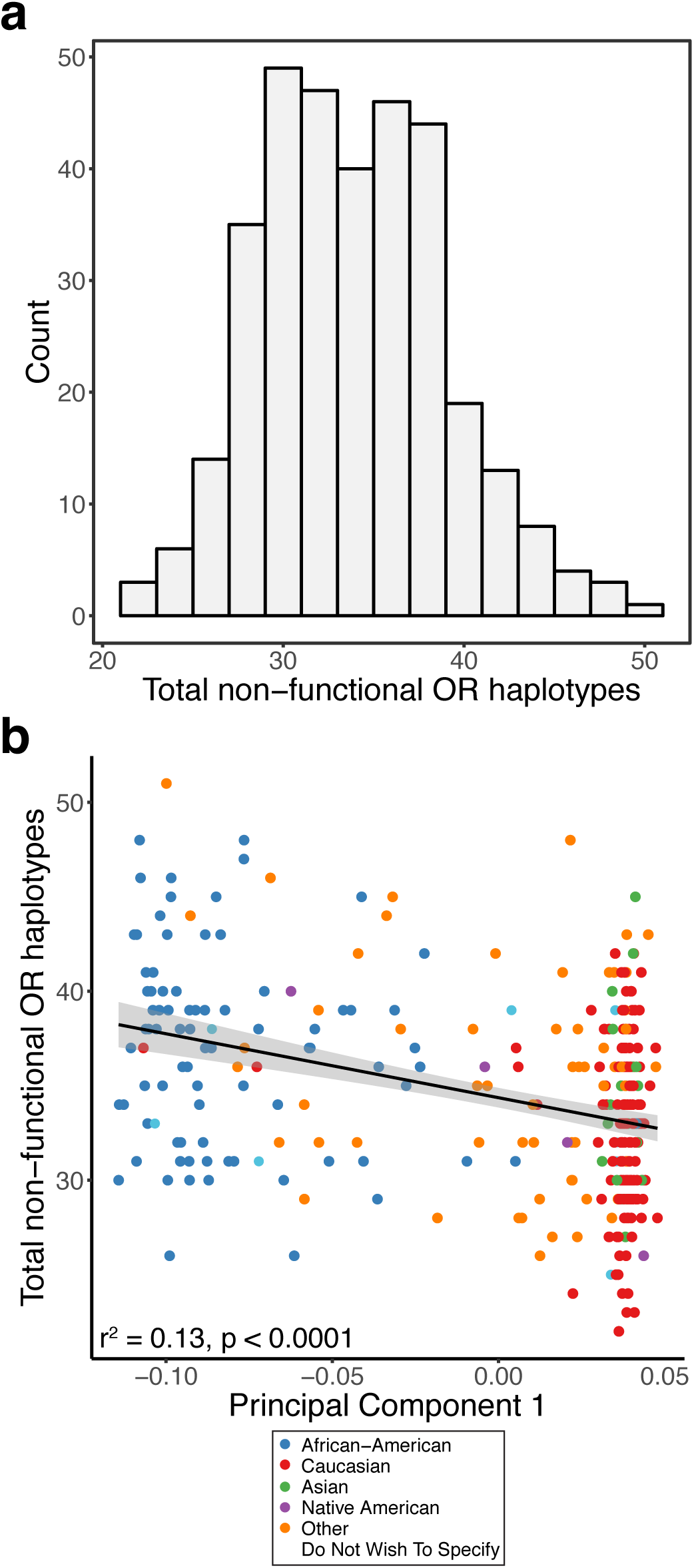
Pseudogene frequency in the participant population. (**a**) Distribution of frequency of OR haplotypes with either nonsense or frameshift mutations. (**b**) Correlation between ancestry (represented using PC1 calculated from all genetic data) and pseudogene frequency (*r*^2^ = 0.13, p < 0.0001).

